# Autoclaving and long-term storage deplete glutamine sources in complex media and affect bacterial phenotypes

**DOI:** 10.1101/2025.10.15.682619

**Authors:** Koichiro Yamada, Kazuya Ishikawa, Kazuyuki Furuta, Shin-Ichi Miyoshi, Makoto Tsunoda, Chikara Kaito

## Abstract

Autoclave sterilization is the most common method for sterilizing reagents and media in biology. However, the effects of heat-induced loss or modification of complex medium components on bacterial growth and phenotypes remain poorly understood. Here, we investigated the impact of autoclaving on glutamine sources in bacterial complex media using an *Escherichia coli* Δ*glnA* mutant, which requires exogenous glutamine for growth. Δ*glnA* exhibited impaired growth in LB medium after autoclaving compared with non-autoclaved LB, whereas its residual growth indicated the presence of heat-stable glutamine sources. Growth assays and HPLC quantification revealed that free glutamine in LB decreased from 58 µM to 12 µM upon autoclaving, while heat-stable glutamine sources remained at 117 µM. Similar growth defects were observed for Δ*glnA* in autoclaved BHI, TSB, and M17 media compared with their non-autoclaved counterparts. Long-term storage of LB at room temperature for 24 weeks also reduced Δ*glnA* growth regardless of autoclaving, compared with freshly prepared LB. Furthermore, supplementation of glutamine sources into glutamine-deficient MHB medium enhanced biofilm formation by *Pseudomonas aeruginosa*. Collectively, these results demonstrate that autoclaving and storage reduce glutamine sources in complex media, thereby influencing bacterial growth and phenotypes.

## Introduction

Glutamine is a critical amino acid for ammonia assimilation in organisms, including bacteria (1). It is synthesized by glutamine synthetase from glutamate and extracellular ammonia and subsequently serves as a nitrogen donor for the biosynthesis of nucleotides and other nitrogenous compounds. In humans, plasma contains 550–750 µM of free glutamine, representing the most abundant free amino acid in circulation (2). Glutamine is particularly important in mammalian cell culture, where increasing glutamine concentrations in medium enhance cell density (3, 4). In bacteria as well, medium glutamine availability affects phenotypes: elevated glutamine enhances the growth of *Streptococcus pneumoniae* and increases antibiotic susceptibility in some methicillin-resistant strains (5, 6).

A distinctive property of glutamine is that it is stable in a dry state but unstable in aqueous solution, where it undergoes hydrolysis of the side-chain amide group to yield glutamate and ammonia, and at elevated temperatures it cyclizes to pyroglutamate (7–9). Because of this instability, glutamine supplementation is often required for rapidly dividing mammalian cells before culture (10).

Autoclave sterilization, which uses pressurized steam above 100°C, is the most widely employed method to sterilize reagents and media in biology, particularly in microbiology. Unlike filtration or chemical sterilization, autoclaving requires no special materials and is cost- and labor-efficient. Complex media ingredients also undergo multiple heat treatments during manufacturing. For example, yeast extract—widely used in bacterial culture—is spray-dried at high temperatures, and beef extract, another amino acid source, is sometimes heated at 90–120°C for an hour during extraction (11). Thus, both media and their components are repeatedly exposed to heat. While heat sterilization of media is routine, several medium constituents, such as vitamin C, B vitamins, hormones, cysteine, and tryptophan, are known to degrade or denature upon autoclaving. In mammalian cell culture, filtration is therefore employed to avoid the inactivation of heat-labile components, including hormones. In contrast, autoclaving is performed routinely for bacterial culture media such as LB. However, because amino acids exist in diverse forms (free and peptide-bound) in complex media, the effects of autoclaving on their stability and bioavailability remain poorly understood.

In this study, we hypothesized that glutamine in complex media is depleted during heat treatments in the manufacturing process and further reduced by autoclave sterilization at the time of preparation, based on its known instability in aqueous solution. Using an *E. coli* Δ*glnA* mutant lacking glutamine synthetase, we demonstrate that autoclaving reduces glutamine sources available to bacteria in complex media. Moreover, supplementation of glutamine into glutamine-deficient Mueller–Hinton broth (MHB) revealed that glutamine depletion suppresses biofilm formation by *P. aeruginosa*.

## Result

### Free glutamine in aqueous solution is markedly reduced by autoclaving

Glutamine is unstable in aqueous solution and decomposes into glutamate and pyroglutamate (7). Previous studies reported that heating glutamine solutions at >100°C for 1 h reduces glutamine to ∼15% of its initial level (9). Based on these findings, we examined the residual level of free glutamine in aqueous solution under standard autoclaving conditions using thin-layer chromatography. After autoclaving at 121°C for 20 or 40 min, no detectable spots were observed (**Fig. 1A and 1B**), indicating that glutamine had decreased to less than 10% of the initial amount. These results confirm that free glutamine in aqueous solution is markedly depleted during autoclave sterilization.

**Fig. 1.**
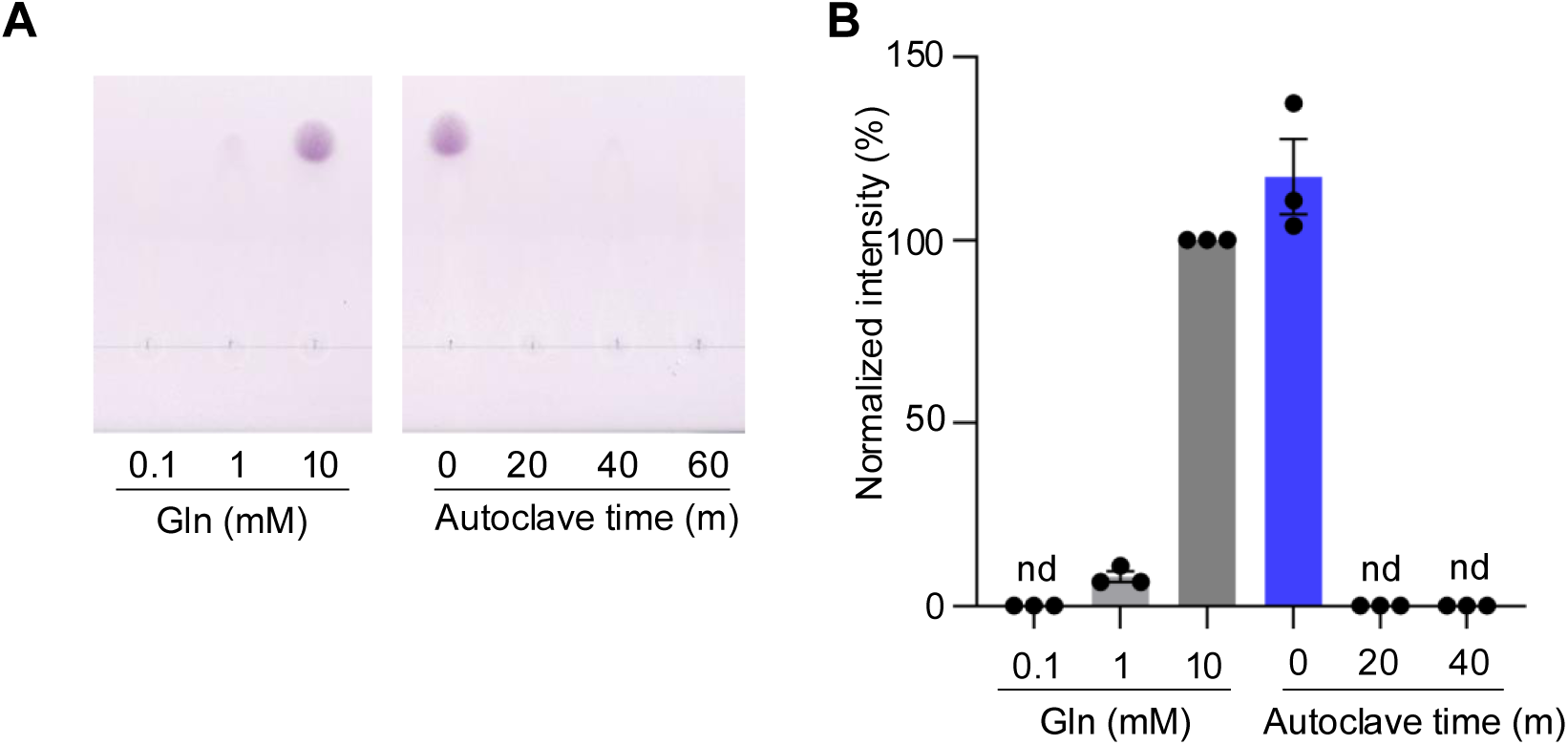
**Free glutamine is markedly reduced by autoclaving.** (A) A 10 mM glutamine solution was autoclaved at 121°C for 20–60 min, separated by TLC, and visualized with ninhydrin. (B) Quantification of glutamine from the TLC signals shown in (A). Values were normalized to 100 using the 10 mM glutamine standard (three independent experiments; mean ± SEM).

### Δ*glnA* exhibits glutamine-dependent growth

To establish a method for estimating glutamine availability in media, we utilized a bacterial strain whose growth depends on exogenous glutamine. We hypothesized that the growth of an E. coli mutant lacking the glutamine synthetase gene (Δ*glnA*) could be used to monitor glutamine concentrations in culture media. To test whether Δ*glnA* grows in a glutamine-dependent manner, we measured its growth in M9 medium supplemented with 0–1000 µM free glutamine. Wild-type cells reached similar stationary-phase OD_595_ values regardless of glutamine supplementation, whereas Δ*glnA* failed to grow without glutamine and exhibited increased stationary-phase OD_595_ values upon glutamine addition (**Fig. 2A and 2B**). Plotting the stationary-phase OD_595_ of Δ*glnA* against the added glutamine concentration revealed a strong positive correlation (R² > 0.98) (**Fig. 2C**). Complementation of Δ*glnA* with a plasmid carrying *glnA* abolished this glutamine requirement (**Fig. 2D**). These results demonstrate that the glutamine-dependent growth of Δ*glnA* is due to the loss of glutamine synthetase activity and indicate that stationary-phase OD_595_ values of Δ*glnA* can be used to estimate glutamine levels in media.

**Fig. 2.**
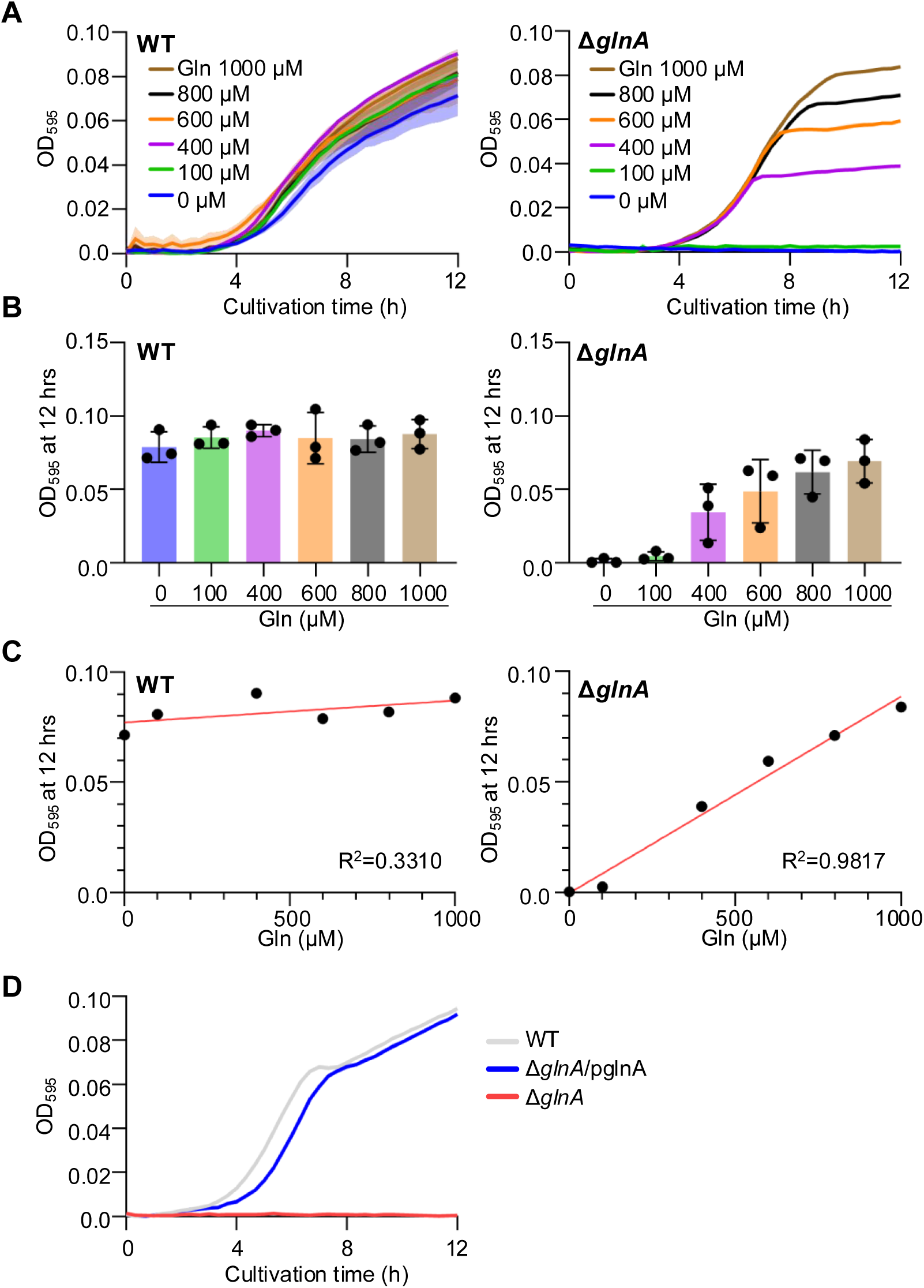
**Δ*glnA* exhibits glutamine-dependent growth.** (A) Growth of WT and Δ*glnA* in M9 minimal medium supplemented with 0–1000 µM glutamine (triplicates; mean ± SEM). (B) OD₅₉₅ values of WT and Δ*glnA* after 12 h of culture in M9 minimal medium with or without glutamine supplementation (three independent experiments; mean ± SEM). (C) Plot of OD₅₉₅ values of WT and Δ*glnA* after 12 h against supplemented glutamine concentration (each point represents the mean of triplicate data). (D) Growth of WT, Δ*glnA*, and complemented Δ*glnA*/pglnA in M9 medium (triplicates; mean ± SEM).

### Glutamine in LB medium is reduced by autoclaving

Because free glutamine was markedly reduced in aqueous solution after autoclaving (**Fig. 1**), we hypothesized that a similar effect occurs in complex media. LB medium, which contains yeast extract and tryptone as amino acid sources, is nutrient-rich but is exposed to heating during both manufacturing and preparation, raising the possibility that glutamine is lost. We therefore evaluated glutamine levels in LB medium using the glutamine-dependent growth of Δ*glnA* as an indicator.

When cultured in autoclaved LB medium, Δ*glnA* exhibited slower growth and markedly lower stationary-phase OD_595_ values compared with the wild type (**Fig. 3A**), suggesting that glutamine was insufficient to support Δ*glnA* growth. To confirm that this growth defect was due to glutamine depletion, we supplemented autoclaved LB with free glutamine. Addition of 100 µM glutamine increased the stationary-phase OD_595_ of Δ*glnA* relative to unsupplemented medium, and 1000 µM supplementation restored the growth curve to a level comparable with the wild type (**Fig. 3B**). These results indicate that the reduced growth of Δ*glnA* in autoclaved LB was caused by glutamine depletion.

**Fig. 3.**
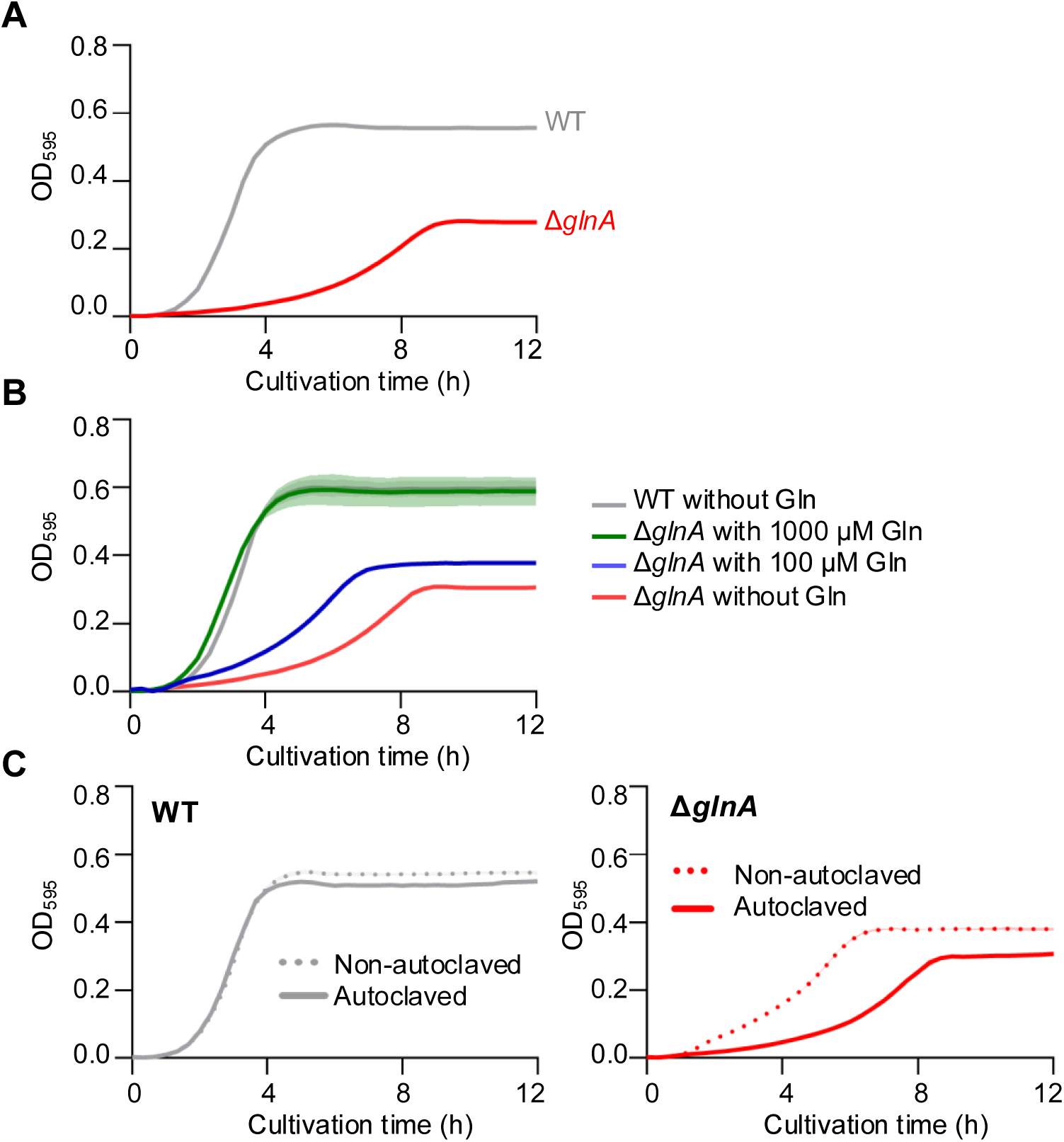
**LB medium is deficient in glutamine.** (A) Growth curves of WT and Δ*glnA* in autoclaved LB medium. (B) Growth curves of Δ*glnA* in autoclaved LB medium supplemented with 100–1000 µM free glutamine. All data are presented as triplicates; mean ± SEM. (C) Growth curves of WT and Δ*glnA* in LB medium sterilized by autoclaving at 121°C for 20 min (solid line) or prepared without autoclaving (dashed line).

To further assess whether glutamine loss occurs specifically during autoclaving, we compared Δ*glnA* growth in LB sterilized by autoclaving versus filter sterilization. Δ*glnA* showed reduced growth in autoclaved LB relative to filter-sterilized LB (**Fig. 3C**), supporting the conclusion that autoclaving reduces glutamine availability.

Finally, to quantify this effect, we measured free amino acids in autoclaved and non-autoclaved LB by high-performance liquid chromatography (HPLC). Free glutamine in autoclaved LB was decreased to approximately one-fifth of that in non-autoclaved LB (**Table 1**). Collectively, these results demonstrate that glutamine in LB medium is substantially depleted by autoclave sterilization.

**Table 1.** Amount of free glutamine in LB medium with or without autoclaving.

|  | **Non-autoclaved LB** | **Autoclaved LB** |
| --- | --- | --- |
| **Amino acid** | **Amino acid concentration (μM)** |  |
| Asn | 832.9 | 1064.1 |
| Asp | 729.8 | 934.5 |
| Gln | 56.1 | 12.6 |
| Glu | 2200.3 | 2717.3 |
| Ser | 1053.4 | 1361.3 |
| Gly | 741.6 | 969.1 |
| His | 341.8 | 444.1 |
| Thr | 934.2 | 1168.5 |
| Ala | 2234.2 | 2765.2 |
| Arg | 1402.8 | 1808.9 |
| Pro | 417.5 | 517.2 |
| Val | 1609.2 | 2030.6 |
| Met | 308.6 | 359.3 |
| Ile | 1175.8 | 1518.1 |
| Leu | 3426.1 | 4277.8 |
| Lys | 2276.6 | 3011.3 |
| Phe | 33.1 | 18.7 |
| Total | 19774.2 | 24978.9 |

### Glutamine depletion occurs in various complex media and their amino acid sources

To determine whether glutamine depletion occurs in complex media other than LB, we compared the growth of Δ*glnA* in autoclaved and non-autoclaved BHI, M17, MHB, NB, and TSB media. In non-autoclaved BHI, M17, and TSB, Δ*glnA* exhibited reduced growth, reaching approximately half the stationary-phase OD_595_ of the wild type (**Fig. 4A**). Growth of Δ*glnA* decreased further when these media were autoclaved, relative to their non-autoclaved counterparts (**Fig. 4A**), indicating that autoclaving exacerbates glutamine depletion in these media. In contrast, Δ*glnA* failed to grow in either autoclaved or non-autoclaved MHB and NB, suggesting that these media contain little or no glutamine.

**Fig. 4.**
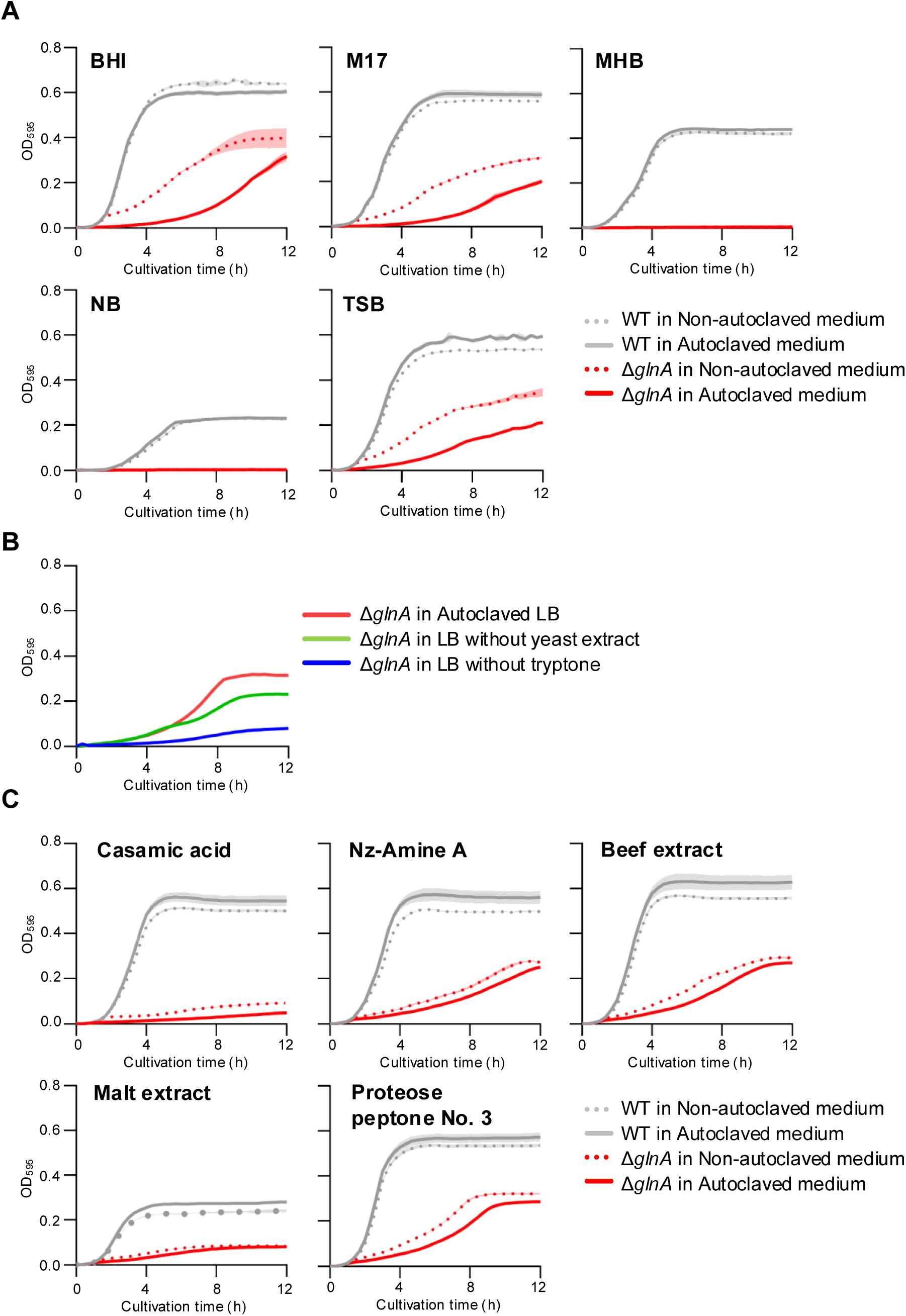
**Autoclaving reduces Δ*glnA* growth in various complex media.** (A) Growth curves of WT and Δ*glnA* in BHI, M17, MHB, NB, and TSB media sterilized by autoclaving (solid line) or prepared without autoclaving (dashed line). (B) Growth curves of WT and Δ*glnA* in LB medium lacking tryptone or yeast extract. (C) Growth curves of WT and Δ*glnA* in LB medium in which tryptone was replaced with Nz-Amine A, beef extract, malt extract, or Proteose peptone No. 3 and sterilized by autoclaving (solid line) or prepared without autoclaving (dashed line). All data are shown as triplicates; mean ± SEM.

We next investigated whether glutamine depletion occurs at the level of the medium components themselves. LB medium consists primarily of yeast extract and tryptone. To assess which component serves as the main glutamine source, we prepared LB lacking either yeast extract or tryptone. Δ*glnA* exhibited reduced growth in LB lacking tryptone compared with LB lacking yeast extract (**Fig. 4B**), indicating that tryptone is the primary glutamine source in LB.

To test whether other amino acid sources exhibit similar autoclave sensitivity, we replaced tryptone in LB with alternative components: beef extract and casamino acids (from MHB medium), Nz-Amine A (an enzymatic hydrolysate of casein), Proteose peptone No. 3 (a milk protein digest), and malt extract. In all cases, Δ*glnA* showed reduced growth compared with the wild type (**Fig. 4C**). Moreover, autoclaving further reduced Δ*glnA* growth in media containing beef extract, casamino acids, Nz-Amine A, or Proteose peptone No. 3, but not in media containing malt extract (**Fig. 4C**). Taken together, these results demonstrate that glutamine depletion occurs not only in LB but also in multiple complex media, and that glutamine sources in commonly used protein hydrolysates—such as tryptone, beef extract, casamino acids, Nz-Amine A, and Proteose peptone No. 3—are diminished by autoclaving.

### LB tryptone contains heat-stable glutamine sources

Although Δ*glnA* showed impaired growth in autoclaved LB medium, it was still able to grow, suggesting the presence of glutamine sources that are resistant to autoclave-mediated degradation. To examine whether prolonged autoclaving would further reduce glutamine availability, we estimated glutamine levels in LB medium after autoclaving for up to 120 min. Compared with 20-min autoclaving, 60-min autoclaving caused a further decrease in Δ*glnA* growth, but the reduction was smaller than that observed between non-autoclaved and 20-min autoclaved LB (**Fig. 5A**). Even after 120 min of autoclaving, glutamine sources were not completely eliminated, indicating the presence of heat-stable glutamine sources in LB.

**Fig. 5.**
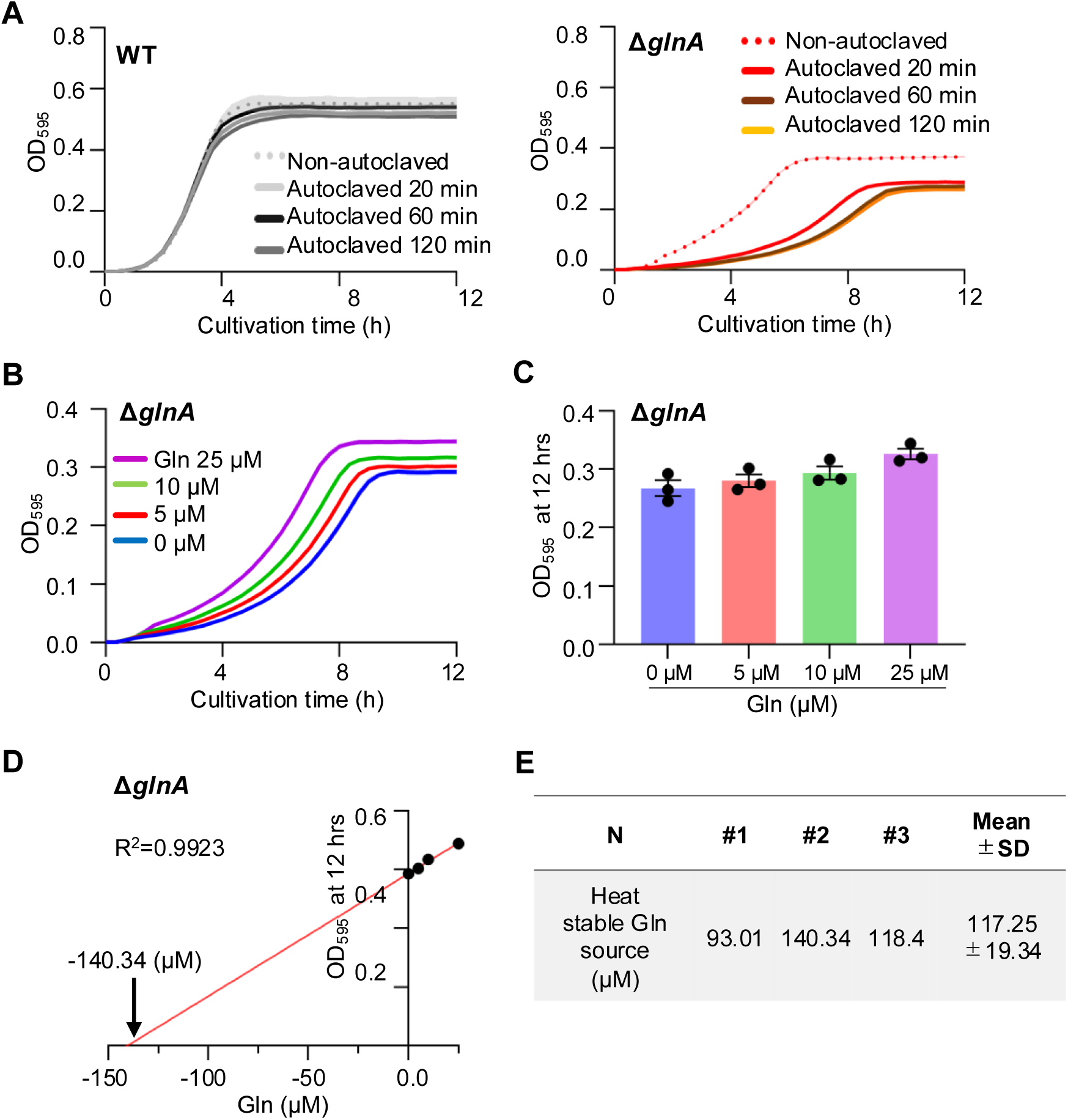
**LB medium contains heat-stable glutamine sources.** (A) Growth curves of WT and Δ*glnA* in LB medium sterilized by filtration or autoclaving at 121°C for 20, 60, or 120 min (triplicates; mean ± SEM). (B) Growth of Δ*glnA* in LB medium supplemented with 0–25 µM glutamine (triplicates; mean ± SEM). (C) OD₅₉₅ values of Δ*glnA* after 12 h in glutamine-supplemented LB medium (three independent experiments; mean ± SEM). (D) Calibration curve constructed by plotting OD₅₉₅ values of Δ*glnA* after 12 h against supplemented glutamine concentrations (each point represents the mean of triplicate data). (E) Biological replicates (n = 3) estimating the concentration of heat-stable glutamine sources in LB medium using the method described in (D).

To quantify these heat-stable glutamine sources, we exploited the glutamine-dependent growth of Δ*glnA*. Similar to the results in M9 medium (**Fig. 2**), supplementation of free glutamine into LB medium enhanced Δ*glnA* growth in a concentration-dependent manner (**Fig. 5B and 5C**). Using this relationship, we generated a calibration curve correlating Δ*glnA* growth with glutamine concentration and applied it to estimate the residual glutamine in autoclaved LB. This analysis indicated that LB medium contains approximately 120 µM of heat-stable glutamine sources (**Fig. 5C and 5D**).

### Long-term storage of LB medium reduces glutamine sources

Glutamine is known to decrease not only upon heating but also over time (8). We therefore examined whether glutamine sources in LB medium decline during prolonged storage. When LB medium was prepared without autoclaving and stored at room temperature, Δ*glnA* growth decreased as the storage period lengthened (**Fig. 6A**). After 24 weeks of storage, OD_595_ values of Δ*glnA* were reduced to 40% at 5 h and 85% at 12 h compared with freshly prepared LB (**Fig. 6A**).

**Fig. 6.**
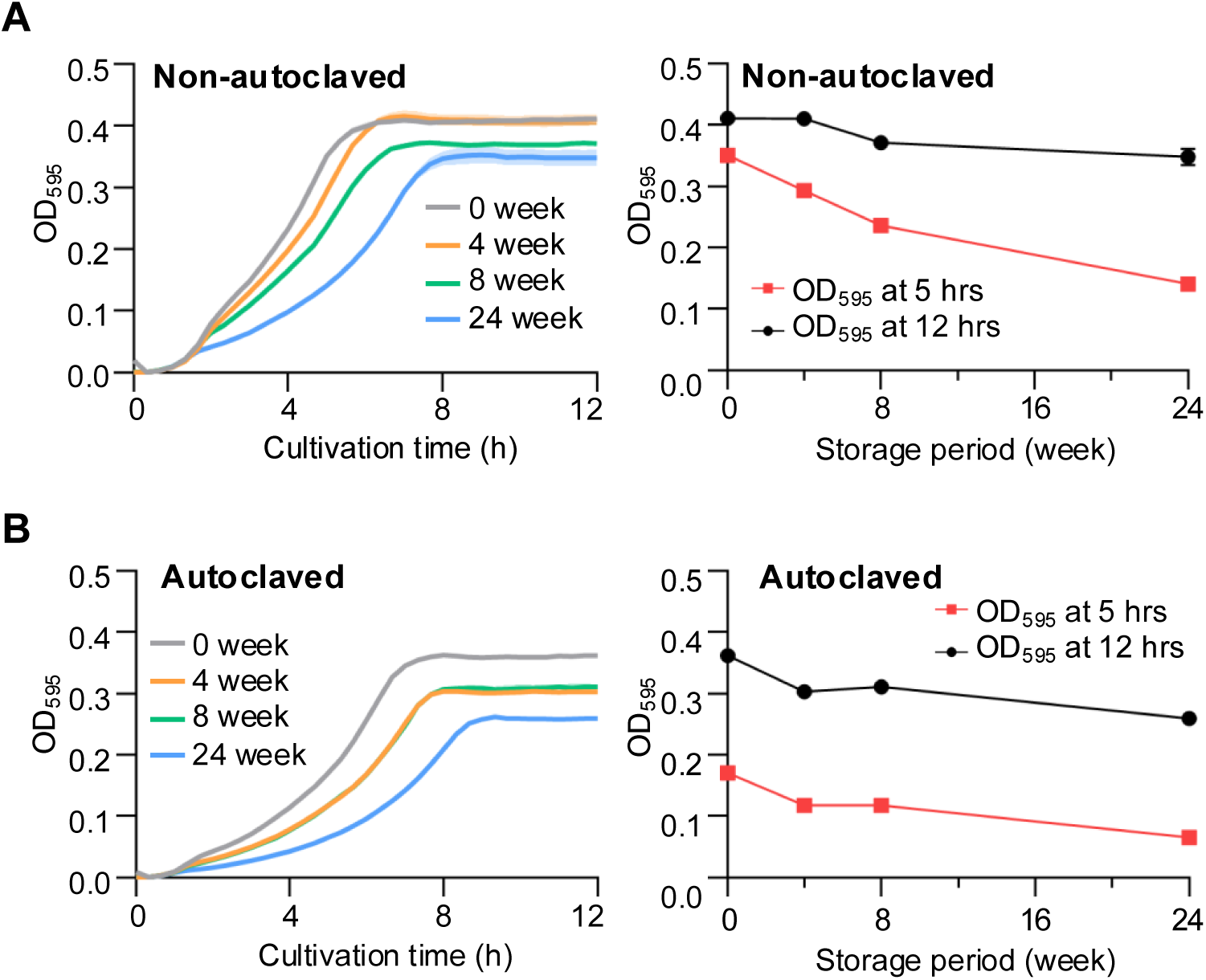
**Long-term storage reduces glutamine sources in LB medium.** (A) Left: Growth curves of Δ*glnA* in LB medium prepared without autoclaving and stored at room temperature for up to 24 weeks. Right: OD₅₉₅ values of ΔglnA at 5 h and 12 h post-inoculation at each storage time point. (B) Left: Growth curves of Δ*glnA* in LB medium prepared by autoclaving and stored at room temperature for up to 24 weeks. Right: OD₅₉₅ values of ΔglnA at 5 h and 12 h post-inoculation at each storage time point. Data represent triplicates; mean ± SEM.

To assess whether heat-stable glutamine sources also diminish over time, we monitored Δ*glnA* growth in autoclaved LB medium stored at room temperature. Δ*glnA* growth decreased progressively with storage duration (**Fig. 6B**). After 24 weeks, OD_595_ values were reduced to 38% at 5 h and 72% at 12 h relative to freshly autoclaved LB (**Fig. 6B**). These findings indicate that both free glutamine and heat-stable glutamine sources in LB medium are reduced during long-term storage at room temperature.

### Glutamine depletion in complex media suppresses biofilm formation by *P. aeruginosa*

Based on the above findings, NB and MHB media appeared to be depleted of glutamine at the level of their raw components, regardless of autoclaving. We next asked whether such glutamine depletion affects bacterial phenotypes. Given that glutamine has been reported to be consumed within *P. aeruginosa* biofilms (12), we examined biofilm formation in glutamine-deficient MHB medium. The amount of biofilm formed by *P. aeruginosa* PAO1 increased significantly when supplemented with ≥750 µM glutamine compared with unsupplemented MHB (**Fig. 7A and 7B**). In contrast, the growth rate of PAO1 was unaffected by glutamine supplementation (**Fig. 7C**). These results suggest that glutamine depletion in complex media suppresses biofilm formation by *P. aeruginosa* PAO1.

**Fig. 7.**
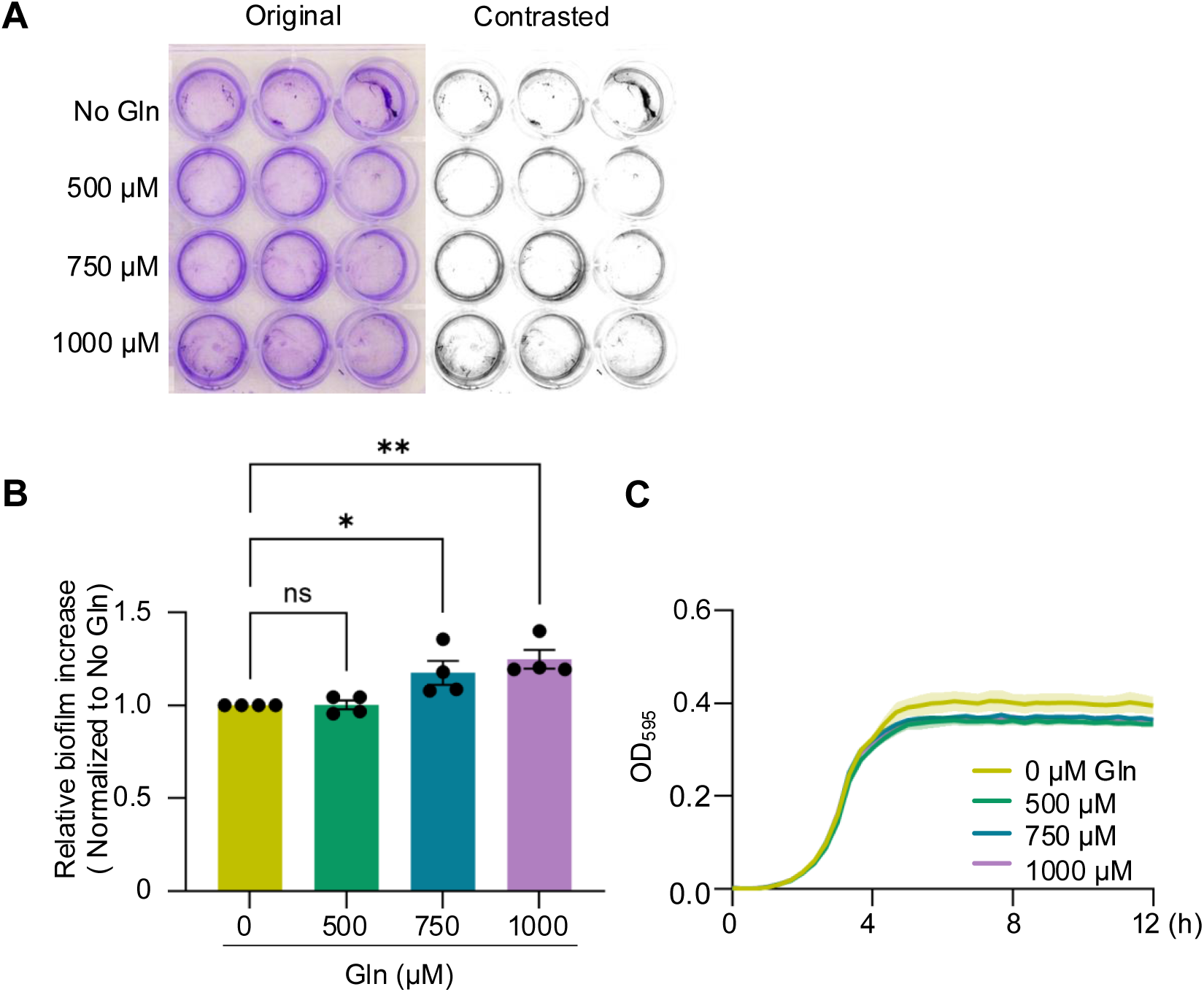
**Supplementation of glutamine enhances biofilm formation by *P. aeruginosa* in MHB medium.** (A) Representative images of biofilms formed by *P. aeruginosa* PAO1 in MHB medium supplemented with 0–1000 µM glutamine, stained with crystal violet. The left image is the original, and the right image is the contrasted version. (B) Quantification of biofilms normalized to the level in the absence of glutamine (set as 1). Data represent more than three independent experiments; mean ± SEM. Statistical significance was determined by one-way ANOVA with Tukey’s post hoc test (*P < 0.05). (C) Growth curves of PAO1 in MHB medium supplemented with 0–1000 µM glutamine (triplicates; mean ± SEM).

## Discussion

In this study, we demonstrated that glutamine sources in complex media, including LB, are depleted by autoclave sterilization and long-term storage, as revealed by the glutamine-dependent growth of an *E. coli* Δ*glnA* mutant. Furthermore, restoration of glutamine availability promoted biofilm formation by *P. aeruginosa*. To our knowledge, this is the first report showing that autoclaving and storage reduce glutamine sources in complex media and thereby influence bacterial phenotypes.

In LB medium, the concentration of free glutamine was already relatively low prior to autoclaving compared with other amino acids (**Table 1**). Although casein-derived tryptone is expected to contain glutamine at levels similar to leucine or glutamate based on amino acid composition (UniProt data), the actual concentration of free glutamine in LB was only 56 µM. Even when heat-stable glutamine sources were included, the total available glutamine remained below 200 µM, suggesting that glutamine is lost during the manufacturing of tryptone.

Glutamine depletion upon autoclaving was also observed in other complex media such as BHI, TSB, and M17 (**Fig. 4A**), which contain different protein hydrolysates as amino acid sources (**Table 2**). Importantly, glutamine reduction occurred in these alternative protein digests as well (**Fig. 4C**), indicating that this effect is not unique to tryptone. A common feature of these amino acid sources is that they are enzymatic digests of proteins, typically prepared under mild conditions (30–55°C for 30 min to 6 h) (13, 14), which likely allows partial preservation of glutamine. By contrast, MHB and NB media contain beef extract and either casamino acids or peptone, respectively. Δ*glnA* failed to grow in either autoclaved or non-autoclaved MHB and NB, suggesting that glutamine had already been lost during production. Beef extract is often prepared by heating beef at >60°C for several hours, and in some cases 90–120°C for 1 h (11), conditions that would degrade glutamine. Casamino acids are produced by acid hydrolysis of casein at 110°C for 24 h in HCl (15), which converts glutamine to glutamate. These production methods likely account for the near-complete depletion of glutamine in MHB and NB.

**Table 2.**
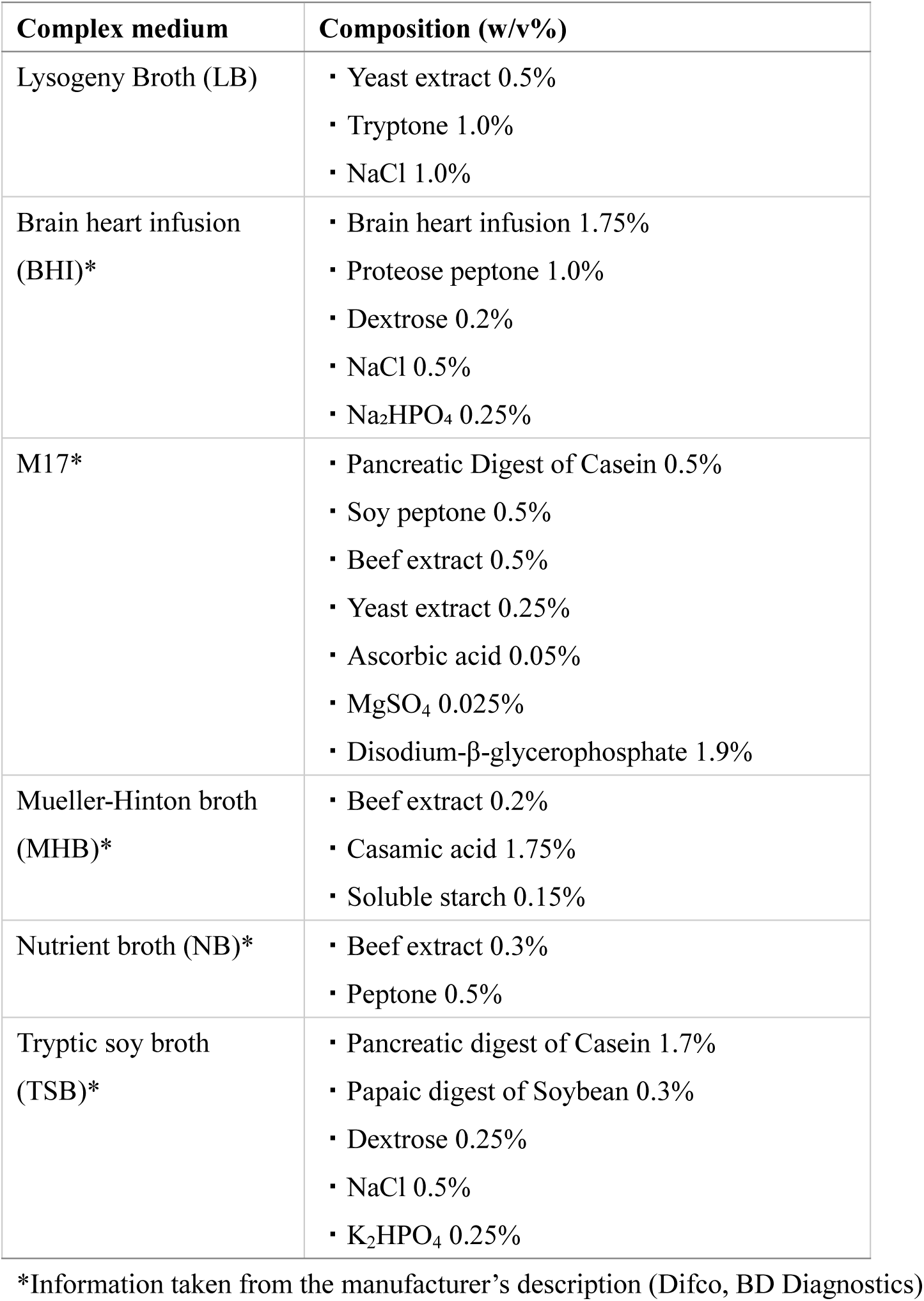
Composition of various complex media.

| Complex medium | Composition (w/v%) |
| --- | --- |
| Lysogeny Broth (LB) | ▪ Yeast extract 0.5%<br>▪ Tryptone 1.0%<br>▪ NaCl 1.0% |
| Brain heart infusion (BHI)* | ▪ Brain heart infusion 1.75%<br>▪ Proteose peptone 1.0%<br>▪ Dextrose 0.2%<br>▪ NaCl 0.5%<br>▪ Na<sub>2</sub>HPO<sub>4</sub> 0.25% |
| M17* | ▪ Pancreatic Digest of Casein 0.5%<br>▪ Soy peptone 0.5%<br>▪ Beef extract 0.5%<br>▪ Yeast extract 0.25%<br>▪ Ascorbic acid 0.05%<br>▪ MgSO<sub>4</sub> 0.025%<br>▪ Disodium-β-glycerophosphate 1.9% |
| Mueller-Hinton broth (MHB)* | ▪ Beef extract 0.2%<br>▪ Casamic acid 1.75%<br>▪ Soluble starch 0.15% |
| Nutrient broth (NB)* | ▪ Beef extract 0.3%<br>▪ Peptone 0.5% |
| Tryptic soy broth (TSB)* | ▪ Pancreatic digest of Casein 1.7%<br>▪ Papaic digest of Soybean 0.3%<br>▪ Dextrose 0.25%<br>▪ NaCl 0.5%<br>▪ K<sub>2</sub>HPO<sub>4</sub> 0.25% |

\*Information taken from the manufacturer's description (Difco, BD Diagnostics)

Our results further suggest that LB contains heat-stable glutamine sources (**Fig. 5**), most likely in the form of peptides. Since amino acids in complex media are predominantly peptide-bound, the heat-stable glutamine fraction is plausibly derived from glutamine-containing peptides. In support of this, dipeptides such as Ala-Gln are known to be more heat-stable than free glutamine and are used as stable glutamine supplements in mammalian cell culture (7). Because tryptone is generated by trypsin digestion rather than acid hydrolysis, it likely retains more peptide-bound glutamine than casamino acids.

We also observed that glutamine levels decrease during long-term storage of LB medium, even in autoclaved preparations enriched in heat-stable glutamine sources (**Fig. 6**). This indicates that although these sources are resistant to heat, they remain unstable over extended periods at room temperature. Thus, both autoclaving and prolonged storage must be considered as factors that reduce glutamine availability in complex media.

Finally, we found that glutamine depletion in complex media suppressed biofilm formation by *P. aeruginosa* PAO1 (**Fig. 7A and 7B**). Given that human plasma contains 550–750 µM free glutamine (2, 16)—42–58 times higher than the free glutamine levels in autoclaved LB and 4–6 times higher even when heat-stable sources are included—our findings raise the possibility that phenotypes observed *in vitro* using complex media may differ from those expressed during infection *in vivo*. Consistent with this, supplementation of ≥1.2 mM glutamine has been reported to enhance biofilm formation in clinical *P. aeruginosa* isolates (17). Together, these findings suggest that glutamine depletion in complex media suppresses biofilm formation and that supplementation restores this phenotype. Beyond biofilm formation, glutamine has been implicated in antibiotic resistance through intracellular pH regulation and in growth control in multiple bacteria (5, 6). Thus, supplementing autoclaved media with glutamine or employing filter sterilization to preserve glutamine may help uncover bacterial phenotypes that have been overlooked due to nutrient depletion during medium preparation.

## Material and Methods

### Bacterial strains and growth conditions

The bacterial strains and plasmids used in this study are listed in Table 3. *E. coli* and *P. aeruginosa* strains were inoculated into 5 mL of antibiotic-free LB medium in 50-mL polypropylene tubes and grown overnight at 37°C with shaking. Overnight cultures were subsequently used for all experiments.

**Table 3.** Strains or plasmids used in this study.

| Strain or plasmid | Genotypes or characteristics<sup>1</sup> | Source of reference |
| --- | --- | --- |
| Strains |  |  |
| *E. coli* |  |  |
| BW25113 | *rrnB*, Δ*lacZ*4787, *HsdR*514, $\Delta$(*araBAD*)567, $\Delta$(*rhaBAD*)568, *rph-1* | NBRP |
| JW3841-KC | BW25113 Δ*glnA::kan* Kan<sup>r</sup> | NBRP |
| JM109 | Host strain for cloning | NBRP |
| *P. aeruginosa* |  |  |
| PAO1 | Laboratory strain | (26) |
| Plasmid |  |  |
| pMW118 | Low-copy-number plasmid; Amp<sup>r</sup> | Nippon Gene |
| pglnA | pMW118 carrying *glnA*; Amp<sup>r</sup> | This study |

<sup>1</sup> Abbreviations: Kan, kanamycin; Amp, ampicillin.

### Thin-layer chromatography (TLC) of glutamine solutions

TLC analysis was performed as previously described (18, 19) with minor modifications. L-glutamine (Wako Pure Chemical Industries, Osaka, Japan) was dissolved in Elix-purified water (Millipore, Billerica, MA, USA) to a final concentration of 10 mM and autoclaved at 121°C for 20, 40, or 60 min. Aliquots (4 µL) of each autoclaved sample, together with L-glutamine standards (0.1–10 mM), were spotted onto silica gel TLC plates (Sigma-Aldrich, St. Louis, MO, USA). Plates were developed for 3 h in a solvent mixture of n-butanol:acetone:diethylamine:Milli-Q water (40:40:3:10, vol/vol). After development, plates were sprayed with ninhydrin reagent (Wako) and heated on a 50°C hot plate to visualize spots. Spot intensities were quantified using ImageJ software.

### Genetic manipulation

Gene deletion by phage transduction was performed as previously described (20, 21). A Δ*glnA*::kan allele from the Keio collection was transduced into *E. coli* BW25113 to generate BW25113 Δ*glnA*::kan (Kan^r^). For complementation, a DNA fragment containing the *glnA* gene was amplified using the following oligonucleotide primers: forward primer (XbaI site), 5’-TCTTCTAGAGTTGGCACAGATTTCGCTTT-3’; reverse primer (HindIII site), 5’-AAGAAGCTTCACATCATCACCGTGTAGGC-3’. The PCR product was cloned into the XbaI and HindIII sites of the pMW118 plasmid to construct the complementation plasmid.

### Preparation of media

M9 medium was prepared by adding 0.6% (wt/vol) KH₂PO₄ (Nacalai Tesque), 0.3% (wt/vol) Na₂HPO₄ (Nacalai Tesque), 0.05% (wt/vol) NH₄Cl (Wako Pure Chemical Industries), 0.1% (wt/vol) NaCl (Nacalai Tesque), 0.4% (wt/vol) glucose (Wako), 2 mM MgSO₄ (Sigma-Aldrich), and 0.1 mM CaCl₂ (Wako) to Elix-purified water (Millipore), followed by sterilization through a 0.22-µm PVDF filter (Millipore).

LB medium was prepared by dissolving yeast extract (Nacalai Tesque), tryptone (Nacalai Tesque), and NaCl (Nacalai Tesque) in Elix-purified water (Millipore) at final concentrations of 0.5% (wt/vol), 1% (wt/vol), and 1% (wt/vol), respectively. Unless otherwise specified, LB medium autoclaved at 121°C for 20 min was used in experiments. Other complex media, including BHI (Difco), M17 (Difco), MHB (Difco), NB (Difco), and TSB (Difco), were prepared according to the manufacturer’s instructions using Elix-purified water (Millipore).

For preparation of LB medium in which tryptone was replaced by alternative amino acid sources, the substitutes were added at 1% (wt/vol), while yeast extract and NaCl were included at the standard LB concentrations. The amino acid sources tested were casamino acids (Difco), Nz-Amine A (Nacalai Tesque), beef extract (Nacalai Tesque), malt extract (Difco), and Proteose peptone No. 3 (Difco). All media were sterilized either by autoclaving at 121°C for 20 min or by filtration through a 0.22-µm PVDF filter (Millipore).

### Measurement of growth curves

Growth curves were measured as described previously (22, 23) with minor modifications. Overnight cultures were centrifuged at 4,000 × g for 5 min at room temperature, and the pellets were washed twice with 0.9% NaCl. When comparing growth between different bacterial strains, OD₆₀₀ values were measured, and the denser culture was diluted with 0.9% NaCl to match the lower OD₆₀₀ value.

Aliquots of 100 µL medium were dispensed into flat-bottom 96-well plates, and 1 µL of washed bacterial suspension was inoculated into each well. Plates were sealed and incubated in a microplate reader (Multiskan FC, Thermo Fisher Scientific, Waltham, MA, USA), which recorded OD₅₉₅ values at 20 min intervals. Each experiment was performed in triplicate wells per sample, and the data are presented as mean ± standard error (SE).

### Glutamine supplementation of M9 medium

L-glutamine was added to M9 medium at final concentrations of 100, 400, 600, 800, and 1000 µM, and the growth curves of the Δ*glnA* strain were measured in these media. OD₅₉₅ values at 12 h post-inoculation were determined in three independent experiments. For each experiment, the mean OD₅₉₅ value of triplicate wells was plotted against the added glutamine concentration. Linear regression analysis was performed using GraphPad Prism 10.

### Quantification of amino acids in culture media by HPLC

Free amino acids in autoclaved or non-autoclaved LB medium were quantified by high-performance liquid chromatography (HPLC). Pre-column derivatization with the amine-reactive reagent 4-fluoro-7-nitro-2,1,3-benzoxadiazole (NBD-F) was performed as described previously (23, 24). Proteins were removed by treatment with 45% methanol/acetonitrile prior to analysis.

### Quantification of heat-stable glutamine in LB medium

Autoclaved LB medium was supplemented with L-glutamine at final concentrations of 0, 5, 10, and 25 µM, and the growth curves of the Δ*glnA* strain were measured in these media. OD₅₉₅ values were determined at 12 h post-inoculation, and the mean values were plotted on the y-axis against the supplemented glutamine concentrations on the x-axis. Linear regression analysis was performed using GraphPad Prism 10, and the absolute value of the x-intercept of the regression line was taken as the concentration of heat-stable glutamine sources in LB medium.

### Long-term storage of LB medium

LB medium prepared by either autoclaving or filtration was transferred into 50-mL polypropylene tubes, wrapped in aluminum foil to protect from light, and stored at 25°C in an incubator. Growth curves of the Δ*glnA* strain were measured immediately after preparation and after 4, 8, and 24 weeks of storage.

### Biofilm assay

Biofilm formation was assessed as described previously (25) with minor modifications. *P. aeruginosa* PAO1 was grown overnight in LB medium, harvested by centrifugation at 4,000 × g for 5 min at room temperature, and washed twice with 0.9% NaCl. MHB medium supplemented with 500–1000 µM L-glutamine (300 µL per well) was dispensed into flat-bottom 24-well plates, and 3 µL of washed culture was inoculated. Plates were incubated statically at 37°C under aerobic conditions for 24 h. Following incubation, the supernatant was removed, and wells were gently washed with 0.9% NaCl to avoid detaching biofilms. Biofilms were stained with 0.1% crystal violet (350 µL per well) for 15 min, washed twice with 0.9% NaCl, and air-dried for 10 min. Crystal violet was then solubilized with 95% ethanol (350 µL per well) for 15 min. Each sample was diluted 10-fold in 0.9% NaCl, and OD₅₉₅ values were measured. Data were normalized relative to the biofilm level in the absence of glutamine (set as 1).

## Acknowledgement

This study was supported by Japan Society for the Promotion of Science (JSPS) Grants-in-Aid for Scientific Research (Grants 23K24131, 23K06130, 24K01760) and the Program of the Japan Initiative for Global Research Network&Link on Infectious Diseases (J-GRID+), JP24wm0125004, from Ministry of Education, Culture, Sports, Science and Technology in Japan (MEXT), and Japan Agency for Medical Research and Development (AMED). We thank the National BioResource Project-*E. coli* (National Institute of Genetics, Japan) for providing the *E. coli* Keio collection.

